# A comprehensive characterization of active expiration in freely behaving rats

**DOI:** 10.1101/2025.02.26.640357

**Authors:** Isabela P. Leirão, Pedro L. Katayama, Daniel B. Zoccal

**Affiliations:** Department of Physiology and Pathology, School of Dentistry of Araraquara (FOAr), São Paulo State University (UNESP), Araraquara, SP, Brazil

**Author notes:** **Corresponding author:** Daniel B. Zoccal, Ph.D. **Address**: Rua Humaitá, 1680, Centro, 14801903; Araraquara, SP, Brazil. Department of Physiology and Pathology, School of Dentistry of Araraquara (FOAR), São Paulo State University (UNESP)., **E-mail:**.

**Keywords:** breathing, active expiration, pulmonary ventilation, hypoxia, hypercapnia

## Abstract

Blood gas disturbances caused by exposure to low oxygen (hypoxia) or high carbon dioxide levels (hypercapnia) lead to a compensatory increase in pulmonary ventilation. Among the motor changes supporting these reflex respiratory responses is the recruitment of abdominal muscles (ABD) during the expiratory phase, which can enhance expiratory airflow or alter the duration of the expiratory phase. In this study, we assessed the functional impact of ABD recruitment on metabolic, motor, and ventilatory parameters in unanesthetized, freely behaving animals. Sprague-Dawley Holtzman male adult rats (n=7) were instrumented to perform simultaneous recordings of pulmonary ventilation, body temperature, diaphragmatic and ABD activities, and O_2_ consumption during exposure to various levels of hypoxia (12-8% O_2_) and hypercapnia (3-7% CO_2_). We observed that hypoxia or hypercapnia conditions evoked AE; however, ABD recruitment did not occur during the entire exposure period, displaying an intermittent profile. The occurrence of AE during hypoxia and hypercapnia conditions was linked to additional increases in tidal volume when compared to periods without ABD activity (P<0.05) and showed no associations with changes in diaphragmatic burst amplitude. Analyses of flow-like patterns suggested that AE during hypoxia recruited expiratory reserve volume during late expiration, while under hypercapnia, it accelerated lung emptying and increased the expiratory flow peak during post-inspiration. We also observed that AE was associated with an increase in oxygen consumption and did not improve air convection requirement, suggesting that this motor behavior may influence other aspects of respiration that potentially improve alveolar ventilation and gas exchange.

**Key points summary:**

- Abdominal recruitment during the expiratory phase, known as active expiration (AE), emerges during blood gas disturbances to enhance pulmonary ventilation; however, the effect of AE on metabolic, motor, and ventilatory parameters in freely behaving animals exposed to hypoxia or hypercapnia remains uncertain.
- By simultaneously recording pulmonary ventilation, diaphragmatic and abdominal activities, and O_2_ consumption, we found that the occurrence of AE during hypoxia or hypercapnia exposure resulted in an additional increase in tidal volume.
- AE was not evident during all periods of exposure to hypoxia or hypercapnia, and its expression increased O_2_ consumption.
- Analysis of flow-like signals suggested that AE during hypoxia recruits expiratory reserve volume in late expiration, while under hypercapnia, it facilitates lung emptying and increases the peak expiratory flow post-inspiration.
- Despite being similar from a motor perspective, the impact of AE on lung ventilation differs between hypoxia and hypercapnia in unanesthetized rats.

## INTRODUCTION

Breathing is essential for maintaining oxygen (O_2_) and carbon dioxide (CO_2_) physiological levels in the arterial blood. The mammalian respiratory motor activity is regulated by a robust yet dynamic neuronal circuitry in the brainstem that coordinates the activity of respiratory muscles controlling airway resistance and thoracic cavity volume (Del Negro *et al*., 2018). Within this network, a group of neurons in the ventral surface of the medulla oblongata forms the so-called pre-Bötzinger complex (preBötC), which intrinsically generates oscillations that determine the respiratory rhythm and drive breathing activity (Smith *et al*., 1991; Tan *et al*., 2008). The oscillatory activity of the preBötC is relayed to pontomedullary premotor and motor neurons that control the pattern (timing, shape, amplitude) of the upper airway, thoracic, and abdominal (ABD) muscles (Richter & Smith, 2014; Del Negro *et al*., 2018). At rest, inspiration initiates with the contractions of muscles (e.g., diaphragm) that expand the thoracic cavity and generate the inspiratory inflow.

Simultaneously, dilator muscles of the upper airways (e.g., genioglossal and posterior cricoarytenoid muscles) contract to reduce airway resistance and facilitate airflow into the lungs (Del Negro *et al*., 2018). With the end of the inspiratory phase, the air is drawn out of the lungs due to inspiratory muscle relaxation and the retraction of the chest and lungs by the elastic recoil forces. Despite being generated passively, the expiratory outflow is regulated during the first stage of expiration (also known as post-inspiration) by the contraction of adductor laryngeal muscles (e.g., lateral cricoarytenoid and thyroarytenoid muscles) that narrow the glottis and reduce the expiratory flow velocity (Dutschmann *et al*., 2014). At the end of the expiratory phase (or stage 2 of expiration), no respiratory muscle activity is observed, and the expiratory flow ceases when the new respiratory cycle begins.

In conditions of elevated metabolic demand or blood gas disturbances, as seen during physical exercise, high altitudes, or in pathological situations promoting hypoxemia and hypercapnia (e.g. sleep-disordered breathing or obstructive lung diseases), the activity of the respiratory network changes to produce compensatory ventilatory responses. In mammals, a reduction in O_2_ or an elevation in CO_2_ levels in the arterial blood triggers an increase in respiratory frequency and tidal volume (Powell *et al*., 1998; Guyenet & Bayliss, 2015). These responses are primarily associated with increased frequency and amplitude of inspiratory muscle contractions driven by the activation of inspiratory premotor and motoneurons in the brainstem and spinal cord, including the preBötC neurons (Barnett *et al*., 2017). Expiration can also become an active process during hypoxemia and hypercapnia, with the presence of rhythmic contractions of the abdominal muscles, especially during the second phase of expiration (Iizuka & Fregosi, 2007; Lemes & Zoccal, 2014). This motor response results from the activation of a conditional expiratory oscillator, located rostral to the preBötC, in the lateral parafacial group (pFL), which connects to premotor expiratory neurons (Janczewski & Feldman, 2006; Abdala *et al*., 2009; de Britto & Moraes, 2016).

Evidence suggests that abdominal muscle contractions during expiration can enhance pulmonary ventilation through different mechanisms. These include accelerating expiratory flow and reducing expiratory time, which facilitates the onset of the next inspiration. Active expiration (AE) can also recruit the expiratory reserve volume to boost tidal volume or promote additional shortening of diaphragmatic muscle fibres during expiration, thus enhancing the muscle tension-contraction relationship and strengthening subsequent inspiration. However, these possibilities can vary according to the species studied (Jenkin & Milsom, 2014) or the stimulus applied, especially considering that hypoxia and hypercapnia can promote distinct time-dependent ventilatory changes (Powell *et al*., 1998) and metabolic adjustments (Frappell *et al*., 1992; Saiki & Mortola, 1996). Moreover, most studies exploring active expiration induced by hypoxia and hypercapnia conditions used high-intensity stimuli, reduced experimental models (*in situ* or *in vitro*), and/or anesthetized animals in which several physiological parameters were artificially maintained steady (e.g., mechanical ventilation, constant body temperature, and intravenous buffer infusion to control blood pH), some reflex mechanisms were eliminated (e.g., vagotomized, paralyzed), and the activity of the CNS was depressed (Janczewski & Feldman, 2006; Abdala *et al*., 2009; Pagliardini *et al*., 2011; Lemes & Zoccal, 2014; Huckstepp *et al*., 2016; Flor *et al*., 2020). While these studies provide relevant evidence regarding the mechanisms of AE, their limitations preclude a comprehensive understanding of the functional impact of forced expiration on pulmonary ventilation under conditions of blood gas disturbances.

In the present study, we simultaneously recorded pulmonary ventilation, diaphragmatic and abdominal muscle activities, and metabolic rate in unanesthetized, freely behaving animals exposed to different levels of O_2_ and CO_2_ to explore the conditions that evoke AE, its correlation with inspiratory motor activity, the influence of AE on ventilatory parameters, and its correlation with the nature (hypoxia vs. hypercapnia) and intensity of stimulus.

## METHODS

### Animals

The experimental protocols and surgical procedures comply with ARRIVE and the Brazilian National Council for Animal Care (CONCEA) guidelines and were approved by the Ethics Committee on the Use of Animals of São Paulo State University, Araraquara, Brazil (Protocol no. 17/2020). All experiments were performed on adult Sprague-Dawley Holtzman male rats (250-300 g) provided by the animal facility of the School of Dentistry of Araraquara (UNESP/FOAR). Animals received food and water *ad libitum* and were housed on a 12 h light−12 h dark cycle (lights on at 7:00 am) in a temperature (23 ± 1°C) and humidity (50±10%) controlled environment. The experiments were performed during the light phase between 9 am and 5 pm.

### Surgical procedures

Seven days prior to the experiments, the animals were anesthetized with a mixture of ketamine (100 mg/kg; i.p.) and xylazine (10 mg/kg; i.p.). After confirming the anesthetic plane, under aseptic conditions, a pair of fine electrodes made from insulated stainless-steel wires were implanted into the diaphragm (DIA) and abdominal muscles (ABD) to record their electrical activity. To access the midcostal DIA, an incision was made at the right superior–lateral portion of the abdomen, following the ribcage border. On the contralateral side, the oblique ABD muscles were identified, and the electrodes were implanted. The electrode loose ends were tunnelled under the skin and attached to an electrical socket on the dorsal region of the animal’s neck. During this surgical procedure, temperature monitoring probes (SubCue Datalogger Standard, Alberta, Canada) were delicately inserted into the animal’s peritoneal cavity through the same incision performed to access the DIA. The temperature probes were programmed to record the animals’ body temperature every 2 minutes, starting 4 hours before the initiation of the experimental protocol until its end.

Incisions were sutured at the end of the procedures, and the animals received antibiotics (penicillin; 30,000 IU, i.m.) and anti-inflammatory medication (ketoprofen; 1 mg/kg, s.c.) and were monitored until they regained consciousness.

### Assessment of pulmonary ventilation and breathing pattern

Pulmonary ventilation was assessed using whole-body plethysmography (Drorbaugh & Fenn, 1955). The animals were kept individually in a closed acrylic chamber (4 L) connected to a gas mixer (supplied with pure O_2_, CO_2_, and N_2_ tanks) and a vacuum pump, allowing continuous ventilation with humidified normoxic/normocapnic, hypoxic/normocapnic, or normoxic/hypercapnic air mixtures (delivered at a flow rate of 1.5 L/min). The chamber also had sensors providing measurements of internal temperature and humidity during the experiments. The breathing-related pressure oscillations inside the chamber were detected by a highly sensitive pressure transducer (Spirometer, ADInstruments, Bella Vista, Australia). These pressure signals were amplified and acquired in a computer using an acquisition system (PowerLab and Labchart 8, ADInstruments, Bella Vista, Australia; sampling rate: 1 KHz). Before starting each experiment, a small animal ventilator (model 683, Harvard Apparatus, USA) was connected to the chamber and used to inject a known air volume (1 ml) at different frequencies (50, 100, 150, and 180 rpm) for volume calibration. From the breathing-related pressure signals, we calculated: i) respiratory frequency (fR; cycles per minute), derived from the time interval between consecutive respiratory peaks; ii) tidal volume (V_T_, mL.100g^-1^), determined from the cycle amplitude, taking into consideration the calibration volume, animal and chamber’s temperature, barometric pressure, and humidity, according to the equation described by (Drorbaugh & Fenn, 1955; Bartlett Jr & Tenney, 1970), and iii) minute ventilation, calculated as the product of fR and V_T_ (mL.100g^-1^.min^-1^). We also analyzed the frequency of sighs (number events/hour), which were identified as augmented breaths (higher than two times baseline V_T_) followed by a prolongation of expiratory time (Li *et al*., 2016).

A first-order derivative was applied to the whole-body plethysmography signals to generate an airflow-like signal (dmV/dt), as previously described (Hernandez *et al*., 2012) and illustrated in Figure 1C. The derivative signals were generated using a 201-point window to optimize the signal-to-noise ratio and used to evaluate the breathing pattern, focusing on the emergence of active expiration. Under resting conditions when expiration is passive, expiratory flow initiates with a rapid increase that peaks during the first half of expiration, or post-inspiration (PI), followed by a slow reduction during the second half of expiration (E2). Active expiration is commonly associated with the emergence of a second peak during the E2 phase, as previously described (Abbott *et al*., 2011; Pagliardini *et al*., 2011). Therefore, we used the derivative signals to identify changes in the inspiratory and expiratory dynamics associated with motor activity changes during exposures to hypoxia or hypercapnia.

**Figure 1.**
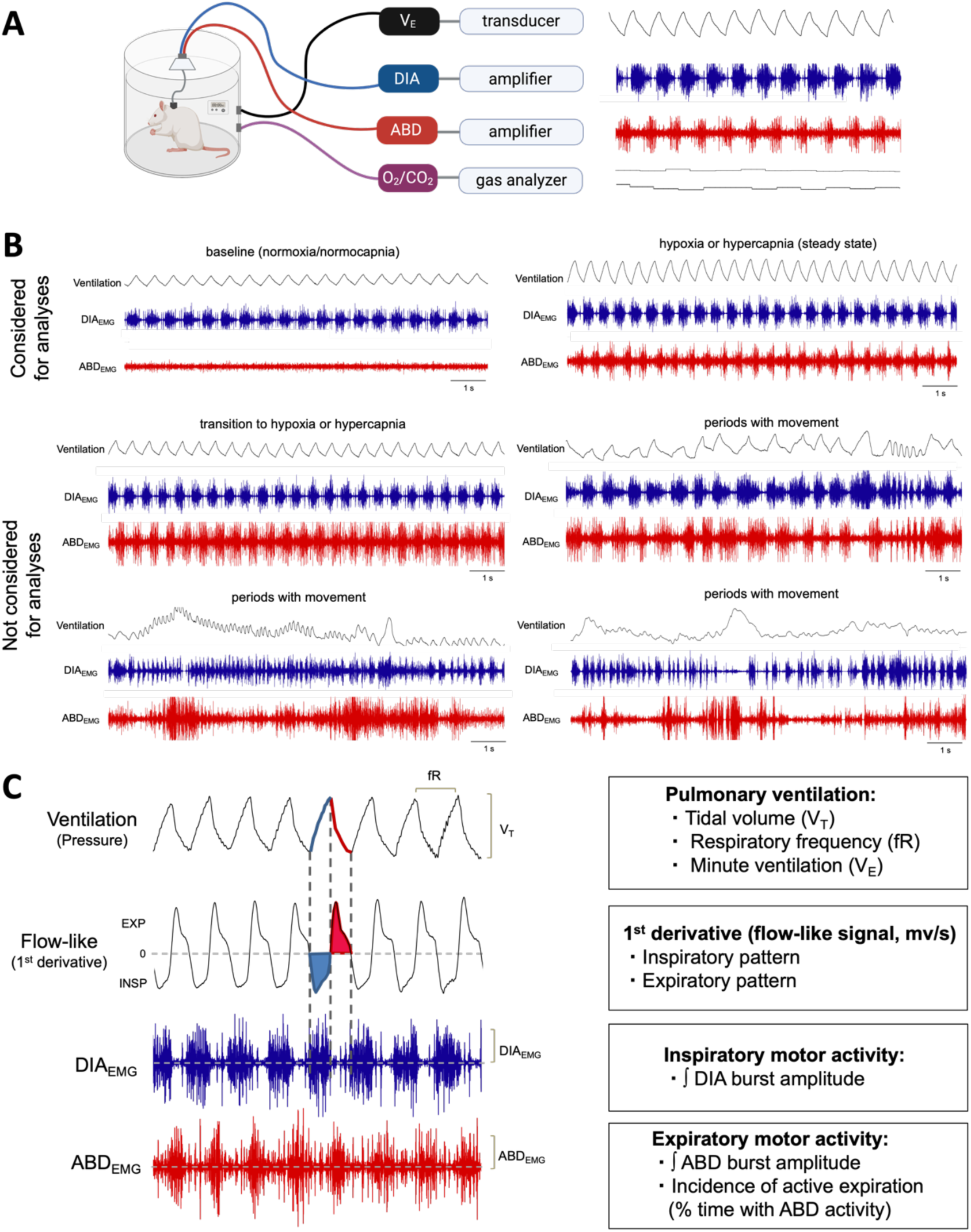
Pulmonary ventilation and EMG recordings in unanesthetized, freely behaving rats. **A.** Schematic diagram and representative traces showing the experimental setup and recorded parameters. Pulmonary ventilation (upper black trace), inspiratory (blue trace) and expiratory (red trace) motor activities were measured in unanesthetized, freely behaving rats (see methods for details). Oxygen and carbon dioxide levels inside the chamber (lower black traces, respectively) were also monitored. **B.** Examples of traces observed during the experiments. Only steady-state periods during baseline, hypoxia, and hypercapnia. The transition periods from baseline to hypoxia or hypercapnia, as well as periods during which the animal moved, were excluded from the analysis. **C.** Methods for data extraction and analyses. From respiratory-related pressure signals (upper trace, black), tidal volume (V_T_) and respiratory frequency (fR) were determined and used to calculate minute ventilation (V_E_). The 1^st^ derivative of ventilation traces generated a flow-like signal (second black trace), allowing the analysis of inspiratory (blue-shaded area) and expiratory (red-shaded area) patterns. The electromyograms of the diaphragm muscle (DIA_EMG_, blue traces) and of the oblique abdominal muscle (ABD_EMG_, red traces) show bursts occurring during inspiration and in the presence of active expiration, respectively.

### O_2_ consumption measurements

Using a flow-through configuration, samples of the inspired and expired gases were drawn through a column of Drierite (Sigma-Aldrich, USA) and then a gas analyzer (ADInstruments, Bella Vista, Australia) to measure the oxygen consumption (VO_2_, mL.100g^-1^.min^-1^) and carbon dioxide production (VCO_2_, mL.100g^-1^.min^-1^), as previously described (Mortola, 1984; Cummings *et al*., 2011). The inflow and outflow airs were sampled continuously, and the values of inspired and expired O_2_/CO_2_ were recorded during all the experiments (PowerLab and LabChart 8, ADInstruments, Bella Vista, Australia; sampling rate: 1 KHz), providing information about the metabolic rate at rest and under hypoxic or hypercapnic conditions. Values of VO_2_ were also used to calculate the air convection requirement (VE/VO_2_).

### Recordings of the diaphragm and abdominal muscle activities in freely behaving animals

The animals were positioned in the plethysmographic chamber to record the electromyographic DIA (DIA_EMG_) and ABD (ABD_EMG_) activities, pulmonary ventilation and metabolic rate under freely behaving conditions. For this purpose, an insulated and shielded cable was connected to the animal’s socket and attached to a swivel on the recording chamber lid. From the lid, a cable was then connected to the amplifier (model P511, Grass Technologies, Middleton, USA) for signal amplification (1000x) and band-pass filtering (10-1000 Hz). EMG signals were acquired in a computer using an acquisition system (PowerLab and LabChart 8, ADInstruments, Bella Vista, Australia; sampling rate: 2 KHz). The raw signals were integrated (smoothed, 0.05 s), and the muscle activity was calculated as the peak amplitude (mV) of the inspiratory and expiratory bursts observed in the integrated DIA_EMG_ (∫DIA_EMG_) and ABD_EMG_ (∫ABD_EMG_), respectively. The AE pattern was defined by the presence of rhythmic ABD_EMG_ activity interposed with DIA_EMG_ activity above tonic levels (Janczewski & Feldman, 2006; Pagliardini *et al*., 2011; Lemes & Zoccal, 2014; Leirao *et al*., 2017; Leirao *et al*., 2020). The incidence of active expiration was calculated as the percentage of time when active expiration was present relative to the total time considered for analyses.

### Experimental protocol

We performed simultaneous recordings of pulmonary ventilation, body temperature, DIA_EMG_ and ABD_EMG_, and O_2_ consumption/CO_2_ production with a minimum of 7 days after the surgical procedures. Animals were placed in a plethysmograph chamber and allowed to acclimate for at least 30 min before starting the experiments. The signals were recorded under resting conditions (normoxia and normocapnia, 30 min) and during exposure to different levels of hypoxia (12%, 10% and 8% O_2_, balanced with N_2_, for 20 min/each) or hypercapnia (3%, 5% and 7% CO_2_, containing 21% O_2_ and balanced with N_2_, for 20 min/each). Each gas condition was tested in the same animal on alternate days. That means that the animals were first exposed to three levels of one gas condition (hypoxia or hypercapnia) on the first day and to the other three levels of the second gas condition (hypercapnia or hypoxia) on the following day. The sequence of conditions and stimulus levels were applied randomly across animals, respecting an interval of at least one hour between consecutive stimuli. After the experiments, the animals were euthanized with anesthesia overdose (sodium thiopental, 200 mg/kg, i.p.) and the temperature probes were removed from the abdominal cavity for data collection and subsequent analyses (The SubCue Analyzer, Alberta, Canada).

### Data and statistical analyses

Analyses were conducted offline using LabChart 8 (ADInstruments, Bella Vista, Australia). As pulmonary ventilation, muscle activity, and metabolic rate were recorded simultaneously, the physiological data were extracted within the same time periods (Figure 1A and C), allowing for correlations between responses. Only periods of quiet breathing and after the stabilization of the gas conditions in the chamber (steady state, 3-5 min after the initiation of exposure) were considered for analyses (Figure 1B). Transitions from normoxia/normocapnia to hypoxia or hypercapnia and periods containing artifacts due to animal movements, exploratory behaviors, and grooming were excluded from the analyses (Figure 1B). All parameters were quantified in their original units and grouped according to the gas condition and intensity. Changes induced by hypoxia or hypercapnia were compared to the corresponding baseline parameters prior to each stimulus. Methods for data extraction and analyses are summarized in Figure 1C.

The results were presented as mean ± SD. The normal distribution of the data was verified using the Shapiro-Wilk normality test. Changes in ventilation, muscle activity, and metabolic rate during hypoxia or hypercapnia were analyzed with one-way ANOVA for repeated measures followed by Tukey’s post-test. Correlations between air convection requirement (V_E_/VO_2_) and active expiration incidence, as well as between VO_2_ and active expiration incidence, were examined using simple linear regressions. Statistical analyses and graphic operations were conducted with GraphPad Prism (GraphPad Software, USA, Version 8). Differences were considered significant when P < 0.05.

## RESULTS

### Active expiration and its functional impact during hypoxia exposure in unanesthetized rats

In our experimental conditions, acute exposure to all hypoxic levels (12%, 10%, and 8% O_2_) increased pulmonary ventilation and respiratory motor activity and reduced metabolic rate (Figure 2 and 3), as previously documented (Frappell *et al*., 1992; Saiki *et al*., 1994). As for the presence of AE, the most intense hypoxic stimulus (8% O_2_) elicited ABD contractions in all animals tested (n = 7), while 10% and 12% O_2_ were able to evoke AE in 6 and 5 (out of 7) rats, respectively (Figure 2A). However, the AE pattern during acute hypoxia was not an all-or-nothing event, as reported in other studies using anesthetized or reduced preparations (Abdala *et al*., 2009; Moraes *et al*., 2012; Lemes & Zoccal, 2014). In fact, we noted that the unanesthetized animals exhibited periods with the presence and absence of AE throughout the period of exposure, with no clear transition pattern. Interestingly, the incidence of AE (i.e., % time the animal presented AE during exposure) was similar across hypoxia levels (Figure 2B).

**Figure 2.**
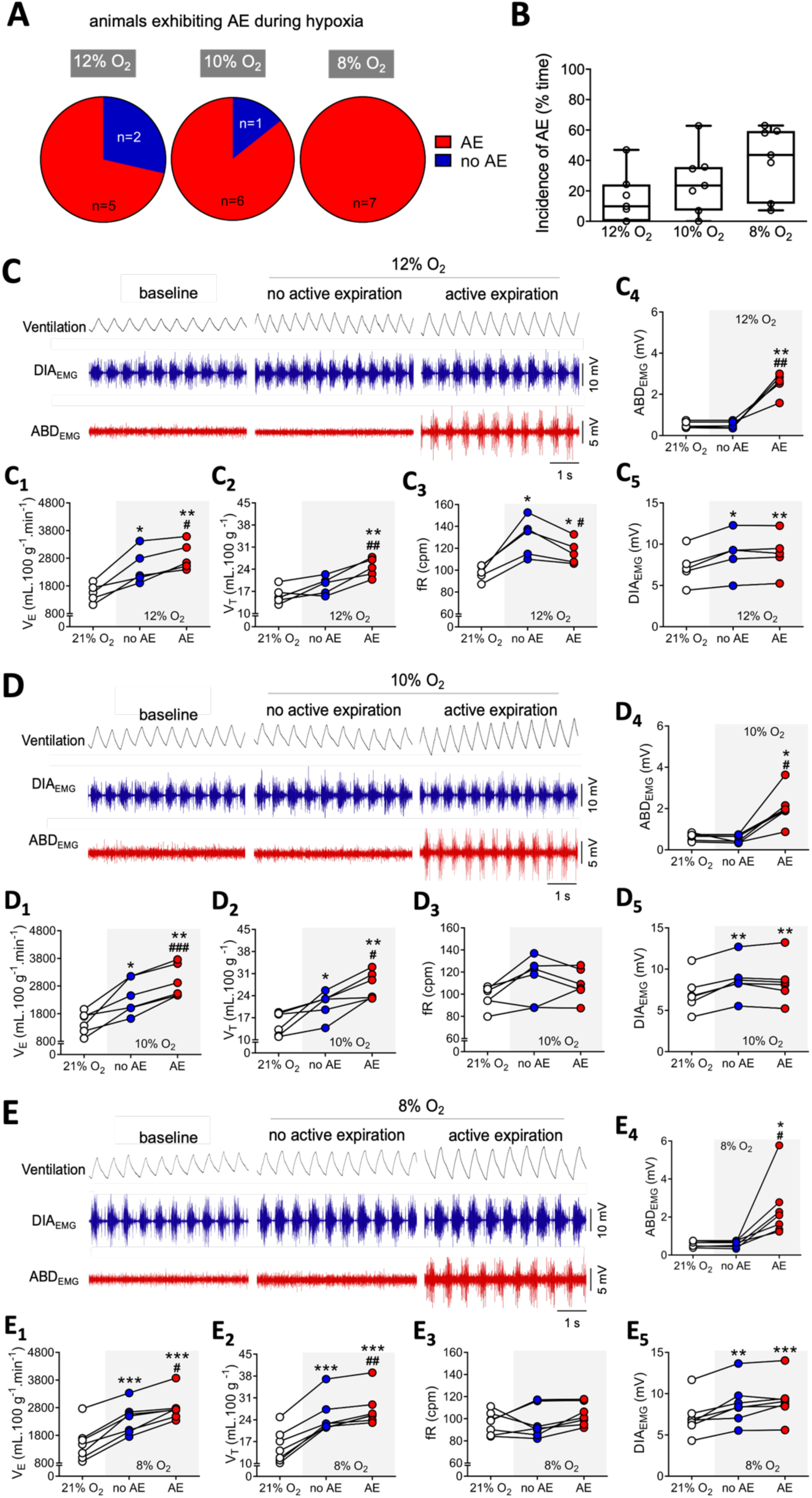
Functional characterization of active expiration during different hypoxia levels. **A.** Pie graphs depicting the proportion of rats presenting (red) or not presenting (blue) active expiration during exposure to different levels of hypoxia (Left, 12% O_2_; middle, 10% O_2_; and right, 8% O_2_). **B.** Boxplots (line represents the mean) showing the incidence of active expiration (% time the animal exhibited ABD muscle recruitment) during exposure to different levels of hypoxia (Left, 12% O_2_; middle, 10% O_2_; and right, 8% O_2_). **C, E and E.** Representative traces of pulmonary ventilation (Ventilation, upper traces, black) and electromyograms of the diaphragm (DIA_EMG_, middle traces, blue) and oblique abdominal muscles (ABD_EMG_, lower traces, red) during baseline (21% O_2_) and hypoxia conditions (C, 12% O_2_; D, 10% O_2_; and E, 8% O_2_). Traces also show periods during hypoxia exposure without active expiration (i.e., absence of ABD rhythmic muscle activity) and with active expiration (i.e., rhythmic ABD muscle activity). **C_1_-C_5_, D_1_-D_5_, and E_1_-E_5_**. Graphs showing individual values of minute ventilation (V_E_; C_1_, D_1_, and E_1_), tidal volume (V_T_; C_2_, D_2_ and E_2_), respiratory frequency (fR; C_3_, D_3_, and E_3_), DIA_EMG_ (C_4_, D_4_, and E_4_), and ABD_EMG_ (C_5_, D_5_, and E_5_) amplitudes, analyzed during baseline periods (baseline, white symbols) and hypoxia periods without (no AE, blue symbols) and with active expiration (AE, red symbols). C_1_-C_5_, 12% O_2_; D_1_-D_5_, 10% O_2_; and E_1_-E_5_, 8% O_2_. For all experiments n=5-7. * P < 0.05 vs 21% O_2,_ ** P < 0.01 vs 21% O_2_, *** P< 0.001 vs 21% O_2_, # P < 0.05 vs no AE, ## P < 0.01 vs no AE, ### P< 0.001 vs no AE (repeated measures one-way ANOVA).

**Figure 3.**
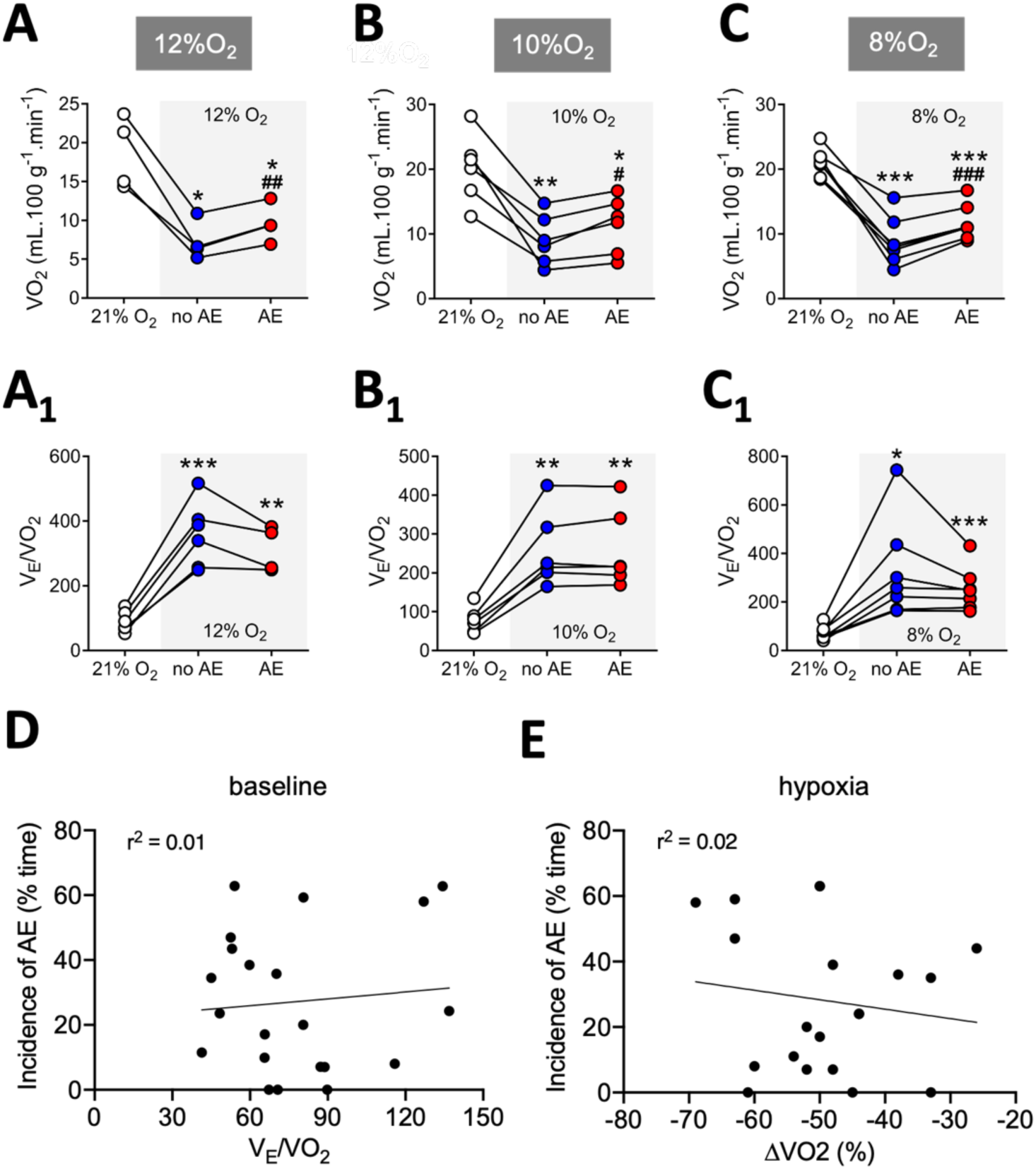
Correlation between active expiration and metabolic measurements during different hypoxia levels. A, B, and. **C.** Graphs showing individual values of oxygen consumption (VO_2_) during baseline (baseline, white symbols) and hypoxia conditions without (no AE, blue symbols) and with active expiration (AE, red symbols). A, 12% O_2_; B, 10% O_2_; and C, 8% O_2_. **A_1_, B_1_, and C_1_.** Individual values of respiratory equivalent (V_E_/VO_2_) during baseline (baseline, white symbols) and hypoxia conditions without (no AE, blue symbols) and with active expiration (AE, red symbols). A_1_, 12% O_2_; B_1_, 10% O_2_; and C_1_, 8% O_2_. **D.** Correlation (linear regression) between the incidence of active expiration (% time the animal exhibited ABD muscle recruitment) and respiratory equivalent (V_E_/VO_2_) during hypoxia. **E.** Correlation (linear regression) between the incidence of active expiration (% time the animal exhibited ABD muscle recruitment) and changes in oxygen consumption (VO_2_) during hypoxia. For all experiments n=5-7. * P < 0.05 vs 21% O_2,_ ** P < 0.01 vs 21% O_2_, *** P< 0.001 vs 21% O_2_, # P < 0.05 vs no AE, ## P < 0.01 vs no AE, ### P< 0.001 vs no AE (repeated measures one-way ANOVA).

To assess the impact of AE on ventilation during hypoxia exposure, we compared, within the same animal, the changes in ventilatory parameters, diaphragmatic activity, and metabolic rate in moments when the ABD expiratory recruitment was present or absent (i.e., no AE) (Figures 2C, 2D, and 2E). In all intensities of hypoxic tested, the emergence of AE promoted a further increase in V_E_ compared to the periods when ABD muscle was not recruited (12% O_2_: 2865 ± 327 vs 2479 ± 622 mL.100g^-1^.min^-1^, P = 0.037, Figure 2C_1_; 10% O_2_: 2958 ± 590 vs 2403 ± 638 mL.100g^-1^.min^-1^, P < 0.001, Figure 2D_1_; 8% O_2_: 2812 ± 495 vs 2388 ± 542 mL.100g^-1^.min^-1^, P = 0.011, Figure 2E_1_). These additional increases in V_E_ were due to higher V_T_ responses in the presence of AE (12% O_2_: 24.5 ± 3 vs 18.9 ± 2.8 mL.100g^-1^, P = 0.009, Figure 2C_2_; 10% O_2_: 27.2 ± 4 vs 21.3 ± 4 mL.100g^-1^, P = 0.018, Figure 2D_2_; 8% O_2_: 27.4 ± 5 vs 24.9 ± 6 mL.100g^-1^, P = 0.005, Figure 2E_2_). The fR was not different from baseline during exposure to 10% and 8% O_2_ (Figures 2D_3_ and 2E_3_). In contrast, the milder hypoxic stimulus (12% O_2_) elevated the fR (no AE vs baseline, 130.2 ± 18 vs 97.7 ± 7 cpm, P = 0.014, Figure 2C_3_) – a response that was attenuated in the presence of AE (116.4 ± 11 vs 130.2 ± 18, cpm, P = 0.04, Figure 2C_3_), but remained above baseline (AE vs baseline, 116.4 ± 11 vs 97.7 ± 7 cpm, P = 0.026, Figure 2C_3_). Hypoxia exposure also strengthened DIA contractions in all levels tested (12% O_2_: no AE vs baseline, 8.8 ± 2.6 vs 7.2 ± 2.1 mV, P = 0.012; AE vs baseline, 8.9 ± 2.5 vs 7.2 ± 2.1 mV, P = 0.003, Figure 2C_5_; 10% O_2_: no AE vs baseline, 8.7 ± 2.3 vs 7.1 ± 2.3 mV, P = 0.001; AE vs baseline, 8.5 ± 2.6 vs 7.1 ± 2.3 mV, P = 0.008, Figure 2D_5_; 8% O_2_: no AE vs baseline, 8.8 ± 2.5 vs 7.1 ± 2.2 mV, P = 0.009; AE vs baseline, 9.2 ± 2.5 vs 7.1 ± 2.2 mV, P < 0.001, Figure 2E_5_). The amplitude of DIA_EMG_ bursts was not altered by the presence of AE (12% O_2_: no AE vs AE, 8.8 ± 2.6 vs 8.9 ± 2.5 mV, P = 0.902, Figure 2C_5_; 10% O_2_: no AE vs AE, 8.7 ± 2.3 vs 8.5 ± 2.6 mV, P = 0.524, Figure 2D_5_; 8% O_2_: no AE vs AE, 8.8 ± 2.5 vs 9.2 ± 2.5 mV, P = 0.296, Figure 2E_5_).

Our analysis of VO_2_ revealed that the hypoxia-induced reduction in the metabolism was smaller during the periods of ABD muscle recruitment (no AE vs AE - 12% O_2_: 9.6 ± 2.4 vs 7.3 ± 2.5 mL.min.100g^-1^, P = 0.009, Figure 3A; 10% O_2_: 11.4 ± 4.3 vs 9.0 ± 3.9 mL.min.100g^-1^, P = 0.016, Figure 3B; 8% O_2_: 11.7 ± 2.7 vs 8.8 ± 3.7 mL.min.100g^-1^, P < 0.001, Figure 3C), indicating that the occurrence of AE was associated with an additional energy expenditure. As a result, the air convection requirement (V_E_/VO_2_), which increased during all hypoxic conditions (P<0.05), remained similar in the absence or presence of AE (no AE vs AE - 12% O_2_: 358.9 ± 100.7 vs 312.7 ± 70.1, P = 0.339, Figure 3A_1_; 10% O_2_: 257.9 ± 96.4 vs 259.4 ± 99.26, P = 0.949, Figure 3B_1_; 8% O_2_: 327.4 ± 205.4 vs 254.1 ± 90.8, P = 0.294, Figure 3C_1_). The incidence of AE did not correlate with baseline V_E_/VO_2_ (r² = 0.01, Figure 3D) or the magnitude of VO_2_ reductions during hypoxia (ΔVO_2_, r² = 0.02, Figure 3E), indicating that neither resting nor hypoxia-induced metabolic responses affected significantly the expression of this motor behaviour.

All levels of hypoxia also increased the frequency of sighs (baseline: 16.7 ± 3.1 events/h vs 12% O_2_: 79.3 ± 29.7 events/h, P = 0.005; 10% O_2_: 80.1 ± 20.1 events/h, P = 0.001; 8% O_2_: 65.0 ± 31.3, P = 0.032, Figures 6A and 6C). The sigh events were frequently followed by a brisk and decrementing ABD burst (Figure 6B). Under hypoxia conditions, when AE was present, we observed a brief period of attenuation of ABD bursts during the post-sigh period (Figure 6B). Based on evidence showing that vagal afferents from lung stretch receptors appear to influence the emergence of AE (Lemes & Zoccal, 2014), we evaluated whether the AE incidence during hypoxia would be influenced by the sigh expression. However, no correlation was found between the incidence of AE and sighs (r^2^ = 0.025, Figure 6D).

### Active expiration and its functional effects during hypercapnia in unanesthetized rats

In conditions of increased inspired CO_2_ levels (3%, 5%, and 7% O_2_), the animals exhibited increased pulmonary ventilation and respiratory motor activity, with no changes in metabolic rate (Figures 4 and 5), as previously reported (Saiki & Mortola, 1996). Similarly to hypoxia, only the most intense hypercapnic stimulus (7% CO_2_) promoted AE in 100% (n = 7) of the animals, while 5% and 3% CO_2_ evoked AE in only 6 and 5 (out of 7) rats, respectively (Figure 4A). Likewise, the AE pattern during acute hypercapnia was not an all-or-nothing event, as described in studies using anesthetized or reduced preparations (Abdala *et al*., 2009; Lemes & Zoccal, 2014), and its incidence was not different across hypercapnic levels (P>0.05, Figure 4B).

**Figure 4.**
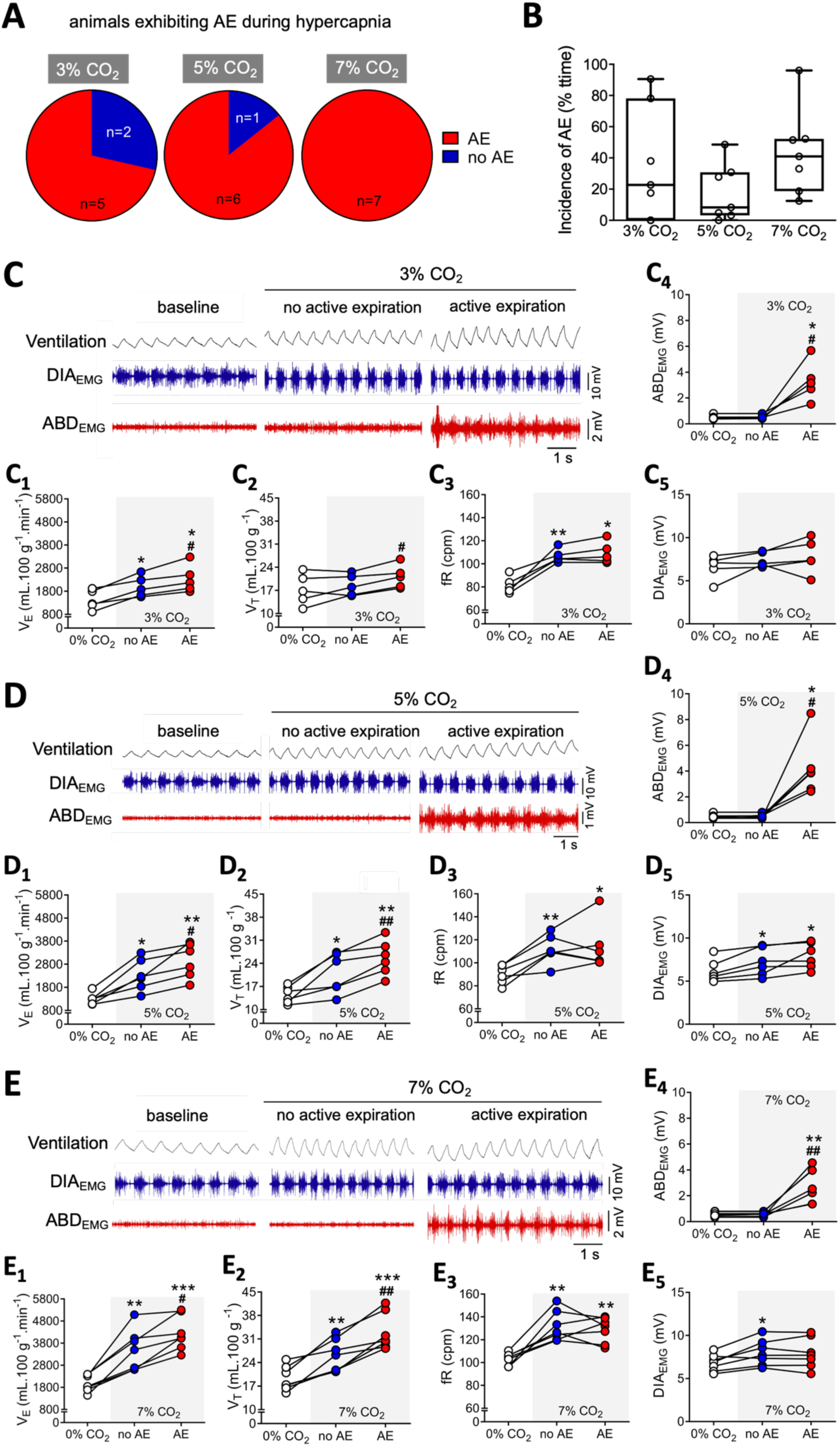
Functional characterization of active expiration during different hypercapnia levels. **A.** Pie graphs depicting the proportion of rats presenting (red) or not presenting (blue) active expiration during exposure to different levels of hypercapnia (Left, 3% CO_2_; middle, 5% CO_2_; and right, 7% CO_2_). **B.** Boxplots (line represents the median) showing the incidence of active expiration (% time the animal exhibited ABD muscle recruitment) during exposure to different levels of hypercapnia (Left, 3% CO_2_; middle, 5% CO_2_; and right, 7% CO_2_). **C, D and E.** Representative traces of pulmonary ventilation (Ventilation, upper traces, black) and electromyograms of the diaphragm (DIA_EMG_, middle traces, blue) and oblique abdominal muscles (ABD_EMG_, lower traces, red) during baseline (0% CO_2_) and during hypercapnia (C, 3% CO_2_; D, 5% CO_2_; and E, 7% CO_2_). Traces also show periods without active expiration (i.e., absence of ABD rhythmic muscle activity) and with active expiration (i.e., rhythmic ABD muscle activity) during hypercapnia exposure. **C_1_-C_5_, D_1_-D_5_, and E_1_-E_5_**. Graphs showing individual values of minute ventilation (V_E_; C_1_, D_1_, and E_1_), tidal volume (V_T_; C_2_, D_2_ and E_2_), respiratory frequency (fR; C_3_, D_3_, and E_3_), DIA_EMG_ (C_4_, D_4_, and E_4_), and ABD_EMG_ (C_5_, D_5_, and E_5_) amplitudes, analyzed during baseline (baseline, white symbols) and hypercapnia conditions without (no AE, blue symbols) and with active expiration (AE, red symbols). C_1_-C_5_, 3% CO_2_; D_1_-D_5_, 5% CO_2_; and E_1_-E_5_, 7% CO_2_. For all experiments n=5-7. * P < 0.05 vs 0% CO_2,_ ** P < 0.01 vs 0% CO_2_, *** P< 0.001 vs 0% CO_2_, # P < 0.05 vs no AE, ## P < 0.01 vs no AE (repeated measures one-way ANOVA).

**Figure 5.**
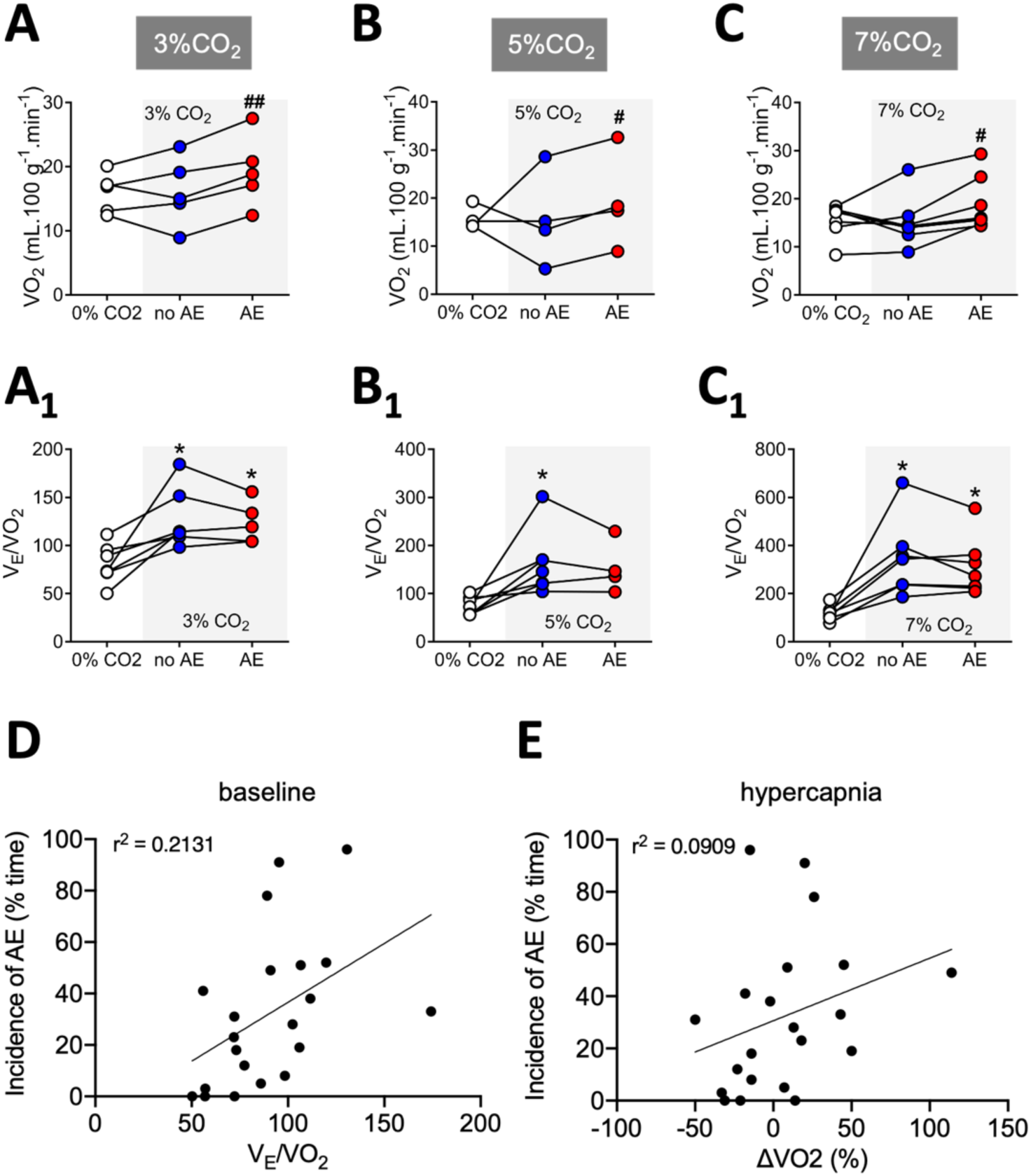
Correlation between active expiration and metabolic measurements during different hypercapnia levels. A, B, and. **C.** Graphs showing individual values of oxygen consumption (VO_2_) during baseline (baseline, white symbols) and hypercapnia conditions without (no AE, blue symbols) and with active expiration (AE, red symbols). A, 3% CO_2_; B, 5% CO_2_; and C, 7% CO_2_. **A_1_, B_1_, and C_1_.** Individual values of respiratory equivalent (V_E_/VO_2_) during baseline (baseline, white symbols) and hypercapnia conditions without (no AE, blue symbols) and with active expiration (AE, red symbols). A_1_, 3% CO_2_; B_1_, 5% CO_2_; and C_1_, 7% CO_2_. **D.** Correlation (linear regression) between the incidence of active expiration (% time the animal exhibited ABD muscle recruitment) and respiratory equivalent (V_E_/VO_2_) during hypercapnia. **E.** Correlation (linear regression) between the incidence of active expiration (% time the animal exhibited ABD muscle recruitment) and changes in oxygen consumption (VO_2_) during hypercapnia. For all experiments n=5-7. * P < 0.05 vs 0% CO_2_, # P < 0.05 vs no AE, ## P < 0.01 vs no AE (repeated measures one-way ANOVA).

As observed in hypoxia, the increases in V_E_ during exposure to all levels of hypercapnia were higher during periods with AE compared to those with no AE (3% CO_2_: 2337 ± 596 vs 1997 ± 455 mL.100g^-1^.min^-1^, P = 0.026, Figure 4C_1_; 5% CO_2_: 2955 ± 758 vs 2334 ± 695 mL.100g^-1^.min^-1^, P < 0.049, Figure 4D_1_; 7% CO_2_: 4247 ± 775 vs 3483 ± 931 mL.100g^-1^.min^-1^, P = 0.020, Figure 4E_1_). These increases in V_E_ associated with AE occurred due to augmented V_T_ responses (3% CO_2_: 21.1 ± 3.5 vs 18.6 ± 3.2 mL.100g^-1^, P = 0.011, Figure 4C_2_; 5% CO_2_: 25.7 ± 5.3 vs 21.0 ± 6.1 mL.100g^-1^, P = 0.001, Figure 4D_2_; 7% CO_2_: 33.0 ± 5.5 vs 26.1 ± 4.8 mL.100g^-1^, P = 0.004, Figure 4E_2_). In contrast, the increase in fR during hypercapnia was not significantly affected by the presence of AE (3% CO_2_: no AE vs baseline, 106.8 ± 9.4 vs 81.6 ± 7.3 cpm, P = 0.009; AE vs baseline, 109.4 ± 9.4 vs 81.6 ± 7.3 cpm, P = 0.012, Figure 4C_3_; 5% CO_2_: no AE vs baseline, 111.8 ± 12.7 vs 89.9 ± 8.2 cpm, P = 0.008; AE vs baseline, 113.9 ± 20.5 vs 89.9 ± 8.2 cpm, P = 0.059, Figure 4D_3_; 7% CO_2_: no AE vs baseline, 131.8 ± 13.2 vs 103.3 ± 5.3 cpm, P = 0.004; AE vs baseline, 128.8 ± 11.0 vs 103.3 ± 5.3 cpm, P = 0.008, Figure 4E_3_). Furthermore, 5 and 7% CO_2_ stimuli, but not 3%, evoked significant increases in DIA_EMG_ burst amplitude (3% CO_2_: no AE vs baseline, 7.4 ± 0.9 vs 6.6 ± 1.4 mV, P = 0.293; AE vs baseline, 7.9 ± 2.0 vs 6.6 ± 1.4 mV, P = 0.099, Figure 4C_5_; 5% CO_2_: no AE vs baseline, 7.2 ± 1.6 vs 6.2 ± 1.3 mV, P = 0.033; AE vs baseline, 8.0 ± 1.5 vs 6.2 ± 1.3 mV, P = 0.011, Figure 4D_5_; 7% CO_2_: no AE vs baseline, 8.0 ± 1.5 vs 6.8 ± 1.0 mV, P = 0.029; AE vs baseline, 7.9 ± 1.7 vs 6.8 ± 1.0 mV, P = 0.105, Figure 4E_5_). The emergence of AE did not modify the hypercapnia-induced changes in the DIA_EMG_ activity (3% CO_2_: no AE vs AE, 7.4 ± 0.9 vs 7.9 ± 2.0 mV, P = 0.783, Figure 4C_5_; 5% CO_2_: no AE vs AE, 7.2 ± 1.6 vs 8.0 ± 1.5 mV, P = 0.261, Figure 4D_5_; 7% O_2_: no AE vs AE, 8.0 ± 1.5 vs 7.9 ± 1.7 mV, P = 0.932, Figure 4E_5_).

Concerning the metabolic rate, the VO_2_ of the animals exposed to hypercapnia remained unchanged compared to baseline values when AE was absent (Figures 5A, 5B, and 5C). On the other hand, VO_2_ was higher in the presence of AE compared to periods without AE (3% CO_2_: 19.3 ± 5.5 vs 16.1 ± 5.3 mL.min.100g^-1^, P = 0.005, Figure 5A; 5% CO_2_: 19.3 ± 9.8 vs 15.6 ± 9.7 mL.min.100g^-1^, P = 0.015, Figure 5B; 7% CO_2_: 19.0 ± 5.7 vs 15.24 ± 5.3 mL.min.100g^-1^, P = 0.015, Figure 5C). Therefore, as observed under hypoxia conditions, the increases in V_E_ associated with AE during hypercapnia (Figures 4C_1_, 4D_1_, and 4E_1_) occurred at the expense of additional O_2_ consumption. As a result, the air convection requirement (V_E_/VO_2_) was not improved during the presence of AE (3% CO_2_: no AE vs AE, 126.4 ± 30.4 vs 123.5 ± 21.8, P = 0.898, Figure 5A_1_; 5% CO_2_: no AE vs AE, 161.8 ± 66.6 vs 153.8 ± 53.6, P = 0.899, Figure 5B_1_; 7% CO_2_: no AE vs AE, 345.3 ± 158.5 vs 311.2 ± 121.3, P = 0.333, Figure 5C_1_). The incidence of AE during hypercapnia did not correlate with baseline V_E_/VO_2_ (r^2^ = 0.021, Figure 5D) or the magnitude of changes in ΔVO_2_ ( ΔVO_2_, r^2^ = 0.09, Figure 5E). Finally, hypercapnia did not change the occurrence of sighs (Figures 6E and 6F); therefore, no association between AE and sigh frequency were noted in this condition.

**Figure 6.**
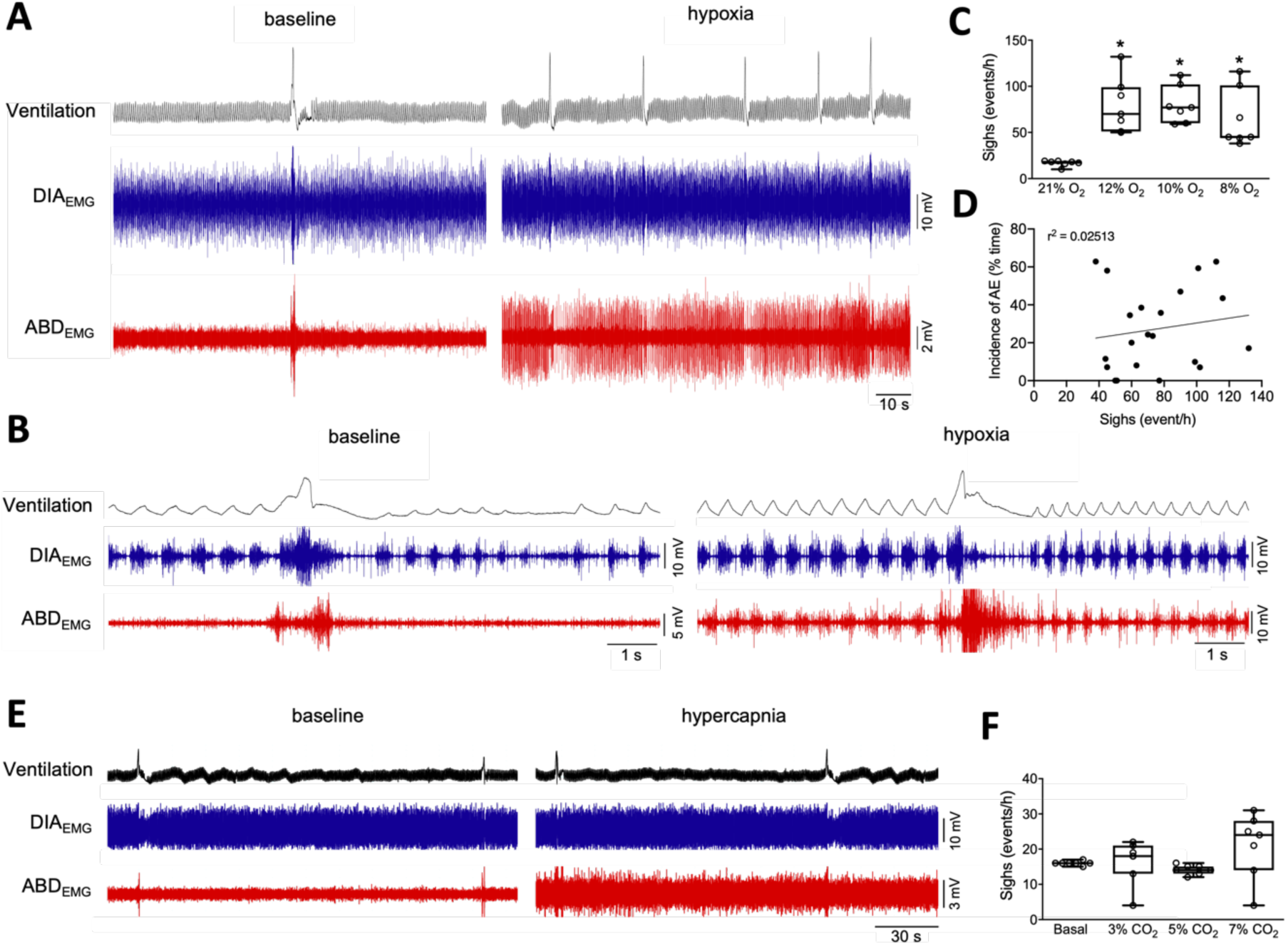
Correlation between active expiration and sighs during hypoxia and hypercapnia. **A.** Representative traces of pulmonary ventilation (Ventilation, upper traces, black) and electromyograms of the diaphragm (DIA_EMG_, middle traces, blue) and of the oblique abdominal muscles (ABD_EMG_, lower traces, red) during baseline (left) and hypoxia conditions (right). **B.** Expanded traces of Ventilation, DIA_EMG_, and ABD_EMG_ showing inspiratory and expiratory motor activity patterns during sighs during baseline (left) and hypoxia (right) conditions. **C.** Occurrence of sighs (events/hour) during baseline (21% O_2_) and different hypoxia levels (12%, 10%, and 8% O_2_). **D.** Correlation (linear regression) between the incidence of active expiration (% time the animal exhibited ABD muscle recruitment) and sigh occurrence (events/hour). **E.** Representative traces of pulmonary ventilation (Ventilation, upper traces, black) and electromyograms of the diaphragm (DIA_EMG_, middle traces, blue) and of the oblique abdominal muscles (ABD_EMG_, lower traces, red) during baseline (left) and hypercapnia conditions (right). **F.** Occurrence of sighs (events/hour) during baseline (0% CO_2_) and different hypercapnia levels (3%, 5%, and 7% CO_2_). * P < 0.05 vs 21% O_2_ (repeated measures one-way ANOVA).

### Effects of active expiration on inspiratory and expiratory dynamics

The airflow-like signals generated by the first-order derivative of whole-body plethysmography signals (dmV/dt) enabled us to estimate the changes in inspiratory and expiratory timing and efforts in animals exposed to hypoxia and hypercapnia and link these changes to the emergence of AE. Using DIA_EMG_ activity as a reference, we divided the respiratory cycle into the following phases: i) inspiration (I), from the onset to the peak of DIA_EMG_ bursts; ii) post-inspiration (PI), coinciding with the decrease in DIA_EMG_ activity after its peak; and iii) stage 2 of expiration (E2), a period marked by the absence of DIA_EMG_ activity. In all animals showing AE under different levels of hypoxia or hypercapnia, we observed that maximal ABD_EMG_ activity occurred during the E2 phase (Figures 7A and 7C). However, the effect on expiratory efforts varied between gas conditions. Under hypoxia (all levels tested), we noted that both inspiratory and expiratory flow-like peaks increased, suggesting enhanced respiratory efforts during both phases (Figure 7B). In the presence of AE (Figure 7B), an additional peak in the expiratory phase during E2 was noted (absent under baseline conditions or without AE), along with a further increase in the inspiratory flow-like peak. During hypercapnic conditions (all levels tested), the inspiratory and expiratory flow-like peaks were higher than baseline, indicating increased respiratory efforts similar to those observed under hypoxia (Figure 7D). However, with AE present, the expiratory flow-like pattern remained unchanged, while significant increases were verified in both the inspiratory and expiratory peaks (Figure 7D). Consequently, these analyses suggest that although ABD_EMG_ reaches its maximal activity during the E2 phase, the influence of AE on airflow and pulmonary mechanics may differ between hypoxia and hypercapnia.

**Figure 7.**
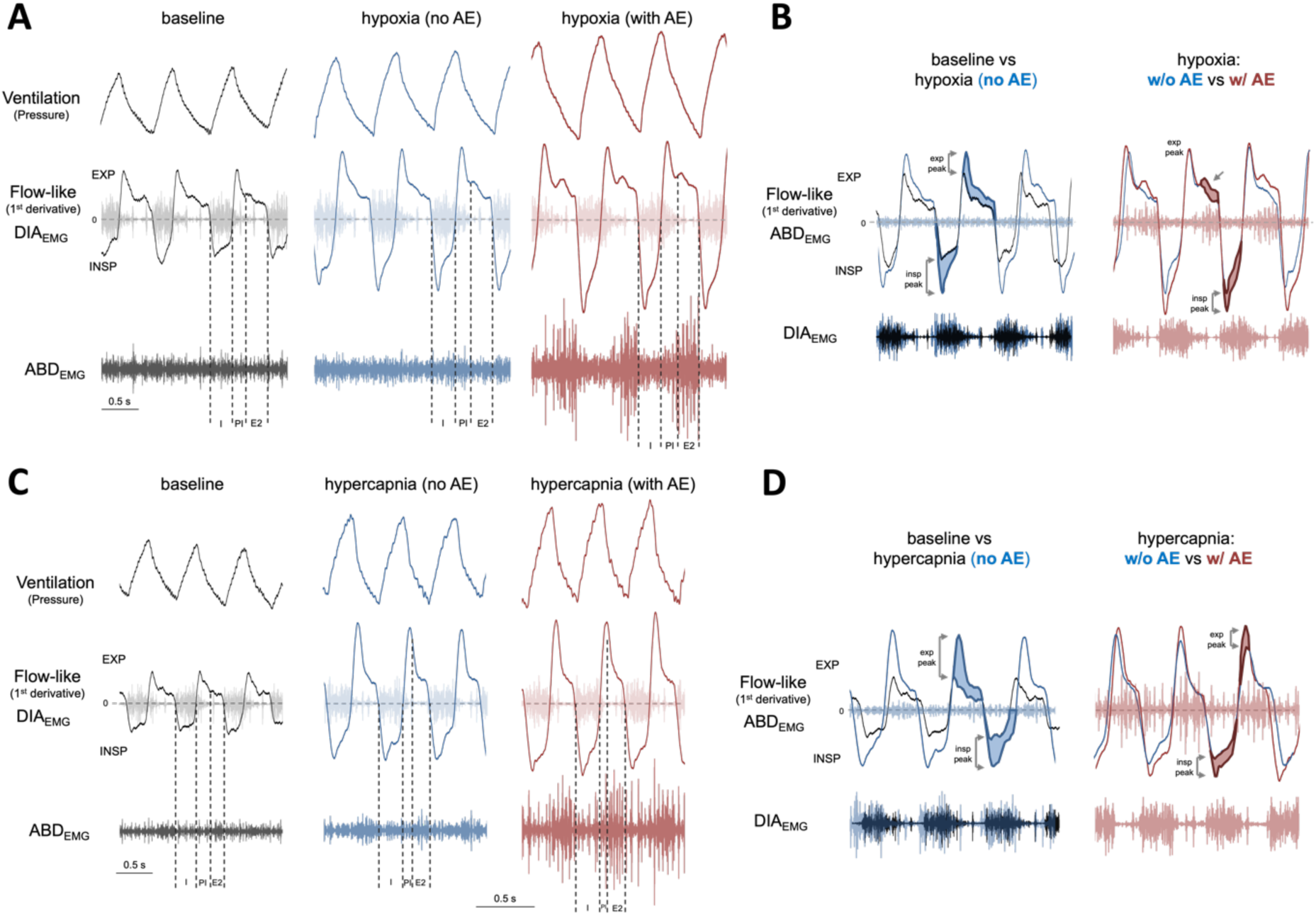
Active expiration and inspiratory/expiratory dynamics during hypoxia and hypercapnia. A and. **C.** Examples of ventilation traces (upper traces), superimposed traces of flow and diaphragm electromyogram (flow-like signal and DIA_EMG_, middle traces), and abdominal electromyogram (ABD_EMG_, lower traces) during baseline (black) and during hypoxia (A) or hypercapnia (C) periods without (blue) and with active expiration (red). Vertical dotted lines indicate respiratory phases: I (inspiration), PI (post-inspiration), E2 (late-expiration). **B and D.** Flow-derived analyses of the inspiratory and expiratory patterns during baseline (black traces) and during hypoxia (B) or hypercapnia (D) periods without (blue traces) and with active expiration (red traces). Inspiratory (insp) and expiratory (exp) peaks are indicated by shaded areas in the flow-like traces.

## DISCUSSION

In mammals, AE is recognized as a respiratory behavior that enhances pulmonary ventilation in situations of blood gas disturbances. A substantial body of evidence in the literature describes the conditions that lead to ABD expiratory recruitment and the corresponding central mechanisms that underpin the emergence of AE pattern (Fregosi & Bartlett, 1988; Mateika *et al*., 1996; Janczewski & Feldman, 2006; Zoccal *et al*., 2008; Abdala *et al*., 2009; Molkov *et al*., 2011; Pagliardini *et al*., 2011; Jenkin & Milsom, 2014; Lemes & Zoccal, 2014; Andrews & Pagliardini, 2015; Burke *et al*., 2015; Huckstepp *et al*., 2015; de Britto & Moraes, 2016; Huckstepp *et al*., 2016; Lemes *et al*., 2016; Jenkin *et al*., 2017; Leirao *et al*., 2017; Barnett *et al*., 2018; Flor *et al*., 2018; Magalhaes *et al*., 2018; Zoccal *et al*., 2018; Flor *et al*., 2020; Leirao *et al*., 2020; Magalhaes *et al*., 2021). Most of these studies were conducted on anesthetized, vagotomized, mechanically ventilated rats or reduced (*in situ*) rodent preparations, which provided detailed and relevant information.

These studies also offered important insights into the potential effects of AE on breathing parameters that would improve pulmonary ventilation. However, limitations in these studies, such as the depressant effects of anesthesia, absence of afferent information, tracheostomy, and clamped metabolic parameters, hindered a comprehensive interpretation of how ABD recruitment during expiration would affect ventilation and blood gas homeostasis. In our study, we combined measurements of pulmonary ventilation, DIA_EMG_ and ABD_EMG_ activities, and metabolic rate to obtain detailed information and advance our comprehension of the ventilatory effects of AE on breathing under conditions of O_2_ deficiency or CO_2_ excess. Our data show that AE evoked during hypoxia and hypercapnia improves minute ventilation mainly by increasing tidal volume without affecting the inspiratory motor activity. However, due to its metabolic cost, AE does not improve air convection requirement (V_E_/VO_2_), suggesting that this motor behavior may have other effects on the respiratory system rather than causing hyperventilation.

The ventilatory responses to hypoxia and hypercapnia are often associated with an increase in the respiratory frequency and diaphragmatic activation caused by the stimulation of inspiratory pre-motor and motoneurons, including the inspiratory rhythm-generating neurons in the preBötC (Zoccal *et al*., 2024). This enhanced inspiratory motor output accelerates breathing rate and increases the depth of inspiration, as seen in our animals exposed to hypoxia and hypercapnia during periods without abdominal recruitment.

Regarding the hypoxic ventilatory response, evidence indicates that it shows a biphasic pattern, with a large initial increase in ventilation, followed by a secondary depression if the hypoxic stimulus persists for several minutes (as in our experiment protocol) mainly due to a reduction in respiratory frequency (Powell *et al*., 1998; Rajani *et al*., 2018). Because our evaluations started 3-5 min after the initiation of the hypoxia exposure, it is likely that the animals were during the secondary phase of the hypoxic ventilatory response, which explains why we did not find relevant changes in respiratory frequency, except at 12% O_2_. We did not consider the initial minutes of hypoxia exposure for analysis because the O_2_ levels in the chamber were still equilibrating, and the animals were frequently moving, which hindered accurate evaluations of EMG activity and O_2_ consumption.

In both hypoxia and hypercapnia situations, regardless of the intensity of the stimulus, the presence of AE amplified pulmonary ventilation due to additional increases in V_T_. These observations support similar observations previously obtained in other experimental conditions (Pagliardini *et al*., 2011; Andrews & Pagliardini, 2015; Huckstepp *et al*., 2015).

However, unlike other studies, we noted that AE did not promote relevant changes in respiratory frequency, except at 12% O_2_. Evidence obtained from *in situ* preparations suggested an association between AE and increased respiratory frequency in response to the activation of O_2_ peripheral chemoreceptors in the carotid bodies (Moraes *et al*., 2012). In contrast, other studies performed in vagotomized and anesthetized animals or reduced preparations showed that AE evoked by hypercapnia or direct activation/disinhibition of expiratory-generating neurons in the pFL leads to expiratory phase prolongation and a reduction in respiratory frequency (Abdala *et al*., 2009; Pagliardini *et al*., 2011; Huckstepp *et al*., 2015; Flor *et al*., 2020). These observations, combined with our results, indicate that AE elicited by hypoxia and hypercapnia primarily affects the lung volume in an unanesthetized animal containing all peripheral afferents and an intact central nervous system. However, we do not exclude that AE may influence respiratory frequency. In another study performed also in unanesthetized conditions, it was reported that AE can be recruited during sleep to promote stability in respiratory frequency when breathing irregularities arise (Andrews & Pagliardini, 2015). We speculate that the effects of AE on respiratory rhythm may depend on the experimental condition or the animal’s state (i.e., sleep or vigilance), which could affect the excitability of the respiratory network.

The changes in V_T_ during periods with AE did not correlate with increases in DIA_EMG_ burst amplitude, suggesting a direct effect of AE on lung volume. The highest ABD activity during hypoxia or hypercapnia occurred in the second stage of expiration (E2). This timing aligns with the firing patterns of expiratory oscillator neurons in the pFL and expiratory motoneurons in the spinal cord (Abdala *et al*., 2009; Moraes *et al*., 2012; de Britto & Moraes, 2017). ABD activation during the E2 phase is thought to enhance expiratory flow and recruit expiratory reserve volume, thereby amplifying the subsequent inspiratory volume (Jenkin & Milsom, 2014). Consistent with this possibility, our analyses of flow-like signals during hypoxia demonstrated that an increased expiratory effort during E2 was associated with an increased inspiratory peak. During hypercapnia, in contrast, the presence of AE was associated with the amplification of expiratory efforts during the first part of expiration (post-inspiration) rather than the E2 phase. These observations suggest that the effect of AE on lung dynamics differs between hypoxia and hypercapnia, even though abdominal muscles are similarly activated during the same phase. We hypothesize that AE increases expiratory flow and recruits expiratory reserve volume during E2 phase during hypoxia exposure, while under hypercapnia, it accelerates lung emptying and augments expiratory flow during the post-inspiratory phase. This difference could happen if these gas conditions differentially modulate upper airway resistance during expiration, restricting expiratory flow during hypoxia and facilitating it under hypercapnia. This possibility aligns with previous reports showing that post-inspiratory laryngeal adductor muscle activity increases during the activation of carotid-body O_2_ chemoreceptors (Barnett *et al*., 2017; Jenkin *et al*., 2017), and that the activity of dilator muscles in the pharynx and larynx increases during hypercapnia (Abdala *et al*., 2009; de Britto & Moraes, 2017). However, these possibilities require further investigation.

A relevant novel finding in our study is the observation that the increase in V_T_ caused by AE does not improve air convection requirement (V_E_/VO_2_). Although this finding may not be surprising - since one might expect an increase in metabolic rate to support muscle contractions - it suggests that the rise in V_E_ induced by AE is proportional to the energy expenditure required for recruiting abdominal muscles, thus not leading to significant improvements in pulmonary ventilation. This interpretation, however, requires additional and detailed experiments to be fully elucidated, including measurements of arterial blood gas during periods with and without AE and analyses of O_2_ extraction ratio. Nevertheless, these findings indicate that the functional significance of AE may not be restricted to alveolar ventilation and may also contribute to other aspects, such as controlling upper airway pressure or distribution of ventilation, which ultimately facilitates gas diffusion, or improving respiratory stability (Mortola *et al*., 1987; Andrews & Pagliardini, 2015). This scenario of various effects of AE on the respiratory system combined with its metabolic cost may explain why this respiratory behavior shows a fragmented and non-uniform expression in unanesthetized animals under hypoxia and hypercapnia. Moreover, evidence shows that the intensity of ABD recruitment can be influenced by the animal’s state or posture (Megirian *et al*., 1987; Sherrey *et al*., 1988; Andrews & Pagliardini, 2015; Leirao *et al*., 2017). Although we believe that the former did not interfere significantly with our analyses due to the short-term nature of our exposure (20 min), the latter is a possibility that was not controlled in our experiments.

In conclusion, our study provides detailed insights into the occurrence and functional impacts of AE on pulmonary ventilation, lung mechanics, respiratory motor activity and metabolism in unanesthetized rats exposed to several levels of hypoxia and hypercapnia. Our data indicate that AE occurs periodically during situations of reduced O_2_ or excessive CO_2_ availability, and enhances expiratory flow and tidal volume to boost lung ventilation independently of increases in diaphragmatic activity. During hypoxia, AE appears to recruit the expiratory reserve volume during the E2 phase while during hypercapnia, AE augments the expiratory flow peak during the post-inspiratory phase to accelerate lung emptying.

However, the onset of AE incurs an additional metabolic cost and does not improve air convection requirement, suggesting that this motor behavior may influence other aspects of the respiratory system that can improve alveolar ventilation and gas exchange. Understanding the physiological role of AE during hypoxia and hypercapnia is crucial, given the prevalence of conditions involving blood gas and metabolic disturbances, such as chronic obstructive pulmonary disease, sleep apnea, or high-altitude acclimatization. Future research exploring the association between impaired AE recruitment and respiratory diseases opens avenues for therapeutic modulation of AE in these contexts.

## AUTHORS’ CONTRIBUTIONS

I.P.L. contributed to conception, design of the work, acquisition, analysis, interpretation of data, drafting and manuscript revision; P.L.K. design of the work, interpretation of data and manuscript revision; D.B.Z. conception, design of the work, analysis, interpretation of data, and manuscript revision. All authors approved the final version of the manuscript and agree to be accountable for all aspects of the work in ensuring that questions related to the accuracy or integrity of any part of the work are appropriately investigated and resolved. All persons designated as authors qualify for authorship, and all those who qualify for authorship are listed.

